# Simple Growth Conditions Improve Targeted Gene Deletion in *Cryptococcus neoformans*

**DOI:** 10.1101/2024.12.18.629221

**Authors:** Rebekah G. Watson, Camaron R. Hole

**Author notes:** Address correspondence to Camaron R. Hole.

## Abstract

*Cryptococcus neoformans* infections are a significant cause of morbidity and mortality among AIDS patients and the third most common invasive fungal infection in organ transplant recipients. The cryptococcal cell wall is very dynamic and can be modulated depending on growth conditions. It was reported that when *C. neoformans* is grown in unbuffered yeast nitrogen base (YNB) for 48 hours, the pH of the media drastically drops and the cells start to shed their cell walls. With this observation, we sought to determine if YNB-grown cells could be used directly for genetic transformation. To test this, we targeted *ADE2* using TRACE (transient CRISPR–Cas9 coupled with electroporation) in YNB-grown or competent cells. Deletion of the *ADE2* gene results in red-pigmented colonies allowing visual confirmation of disruption. We were able to successfully delete *ADE2* in YNB-grown cells and with better efficiency compared to competent cells. Recently, it was shown that gene deletion can be accomplished using short (50 bp) homology arms in place of the normal long arms (∼1 kb). However, it was inefficient, leading to more insertions and gene disruption than gene deletions. We tested short homology with YNB-grown cells vs. competent cells and found gene deletion was significantly improved in YNB-grown cells, around 60%, compared to around 6% with the competent cells. This was also observed when we deleted LAC1 with the short arms. Altogether, using simple growth conditions, we have greatly improved the speed and efficiency of cryptococcal genetic transformations.

**Importance:** The World Health Organization recently ranked *C. neoformans* as the #1 highest-priority fungal pathogen based on unmet research and development needs and public health importance. Understanding cryptococcal pathogenicity is key for developing treatments. We found that using simple growth conditions can greatly improve the speed and efficiency of cryptococcal genetic transformations. This finding will advance the field by expanding the ease of cryptococcal genetic manipulations.

## Introduction

*Cryptococcus neoformans* is a ubiquitous encapsulated fungal pathogen that causes pneumonia and meningitis in immunocompromised individuals. *C. neoformans* infections are a significant cause of morbidity and mortality among AIDS patients and the third most common invasive fungal infection in organ transplant recipients (1–3). Understanding cryptococcal pathogenicity is key for developing treatments. *Cryptococcus* is genetically amenable, and the ability to manipulate *Cryptococcus* genetically has advanced our understanding of the organism.

There are multiple methods used to manipulate *Cryptococcus* genetically. *Agrobacterium tumefaciens*-mediated transformation (ATMT) has been used to make cryptococcal insertion mutagenesis libraries (4). However, targeted gene disruptions are not possible with ATMT, as the T-DNA randomly inserts into the cryptococcal genome (4). Until recently, biolistic transformation was the standard method of cryptococcal genetic manipulation. This method has been used for years to delete and manipulate genes in *Cryptococcus* successfully; however, it has some problems. Besides the high cost of the biolistic system and the consumables, the number of transformants can vary, and a high number of transformants may have to be screened to find the desirable mutant due to low rates of homologous recombination (HR) (5). Another method used is electroporation. However, while electroporation has high transformation efficiency, it was rarely used as it leads to mainly episomally maintained DNA (6). Additionally, the Clustered regularly interspaced short palindromic repeats (CRISPR)-associated protein 9 (Cas9) system has been developed for cryptococcal genetic manipulation (5, 7, 8) 27503169). CRISPR-Cas9 creates a double-stranded break (DSB) at a target location, which helps facilitate donor DNA integration at the DBS by HR repair. The original cryptococcal CRISPR-Cas9 systems required the integration of Cas9 into the genome (7) or an auxotrophic strain (8). These issues were overcome by developing the TRACE (transient CRISPR–Cas9 coupled with electroporation) system. In this system, a Cas9 expression cassette, a target-specific sgRNA expression cassette, and donor DNA are mixed and introduced into the cryptococcal cells by electroporation (5, 9). After HR repair, the Cas9 and sgRNA expression cassettes are eliminated without integration into the genome. TRACE is efficient and versatile and has surpassed biolistic transformation as the standard method of cryptococcal genetic manipulation. This system was further improved by using *Cryptococcus*-optimized Cas9 (10). Multiple groups are actively working to expand this system.

The cryptococcal cell wall is dynamic and can be modulated depending on growth conditions. Depending on the media, temperature, and pH, we can artificially induce changes in cell morphology, cell size, and cell wall composition (11). Most fungal cell walls contain chitin. However, the cryptococcal cell wall is unusual because the chitin is predominantly deacetylated to chitosan. Recently, it was found that changing the media and pH can drastically change the levels of chitosan in the cell wall (12). Cryptococcal cells grown in unbuffered Yeast Nitrogen Base (YNB) media for 48 hours led to significant changes in the capsule, cell wall organization, and a decrease in chitosan (12). Cells grown in unbuffered YNB also had significant cell wall damage that was reversed by shifting the cells to more alkaline and nutrient-rich media, highlighting how growth conditions can drastically change the fungal cell wall (12).

Treatments that degrade or make the fungal cell wall more permeable can improve transformation efficiency. Since the cryptococcal cells grown in unbuffered YNB had significant cell wall damage, we questioned if YNB-grown cells could be used directly for genetic transformation by electroporation. We found that YNB-grown cells can be used directly for genetic transformation and are more efficient than traditional competent cells with both long and short homology deletion constructs. Additionally, YNB-grown cells increased the number of gene deletions over gene disruptions when using short homology arms. By using simple growth conditions, we have greatly improved the speed and efficiency of cryptococcal genetic transformations.

## Results

### YNB-grown cells can be used directly for genetic transformation

Recently, it was reported that growing cryptococcal cells in unbuffered Yeast Nitrogen Base (YNB) media for 48 hours led to significant changes in the capsule and cell wall organization observed by electron microscopy (12). These YNB-grown cells had regions where part of the capsule and outer wall were stripped away from the cells, exposing the inner wall fibrils and a less electron-dense plasma membrane (12). With this observation, we sought to determine if YNB-grown cells could be used directly for genetic transformation by electroporation. Traditionally, to make cryptococcal cells electrocompetent, cells freshly grown from glycerol stocks are incubated overnight in YPD media, back diluted, and grown until the culture reaches the mid-log phase. The cells are washed with water, resuspended in an osmotically supportive media, and incubated on ice with dithiothreitol (DTT) for one hour. The cells are then washed to remove the DTT (Figure 1a). Adding the time it takes to make the cells competent, in addition to the transformation, recovery, and plating time, takes around 9-10 hours to complete. Since the YNB-grown cells have exposed inner cell walls, potentially, these cells would not have to be made competent, leading to a significantly shorter transformation day. To test this, we grew cryptococcal cells in YNB media for 48 hours, washed them with water to remove salts, and resuspended them in an electroporation buffer (Figure 1b).

**Figure 1:**
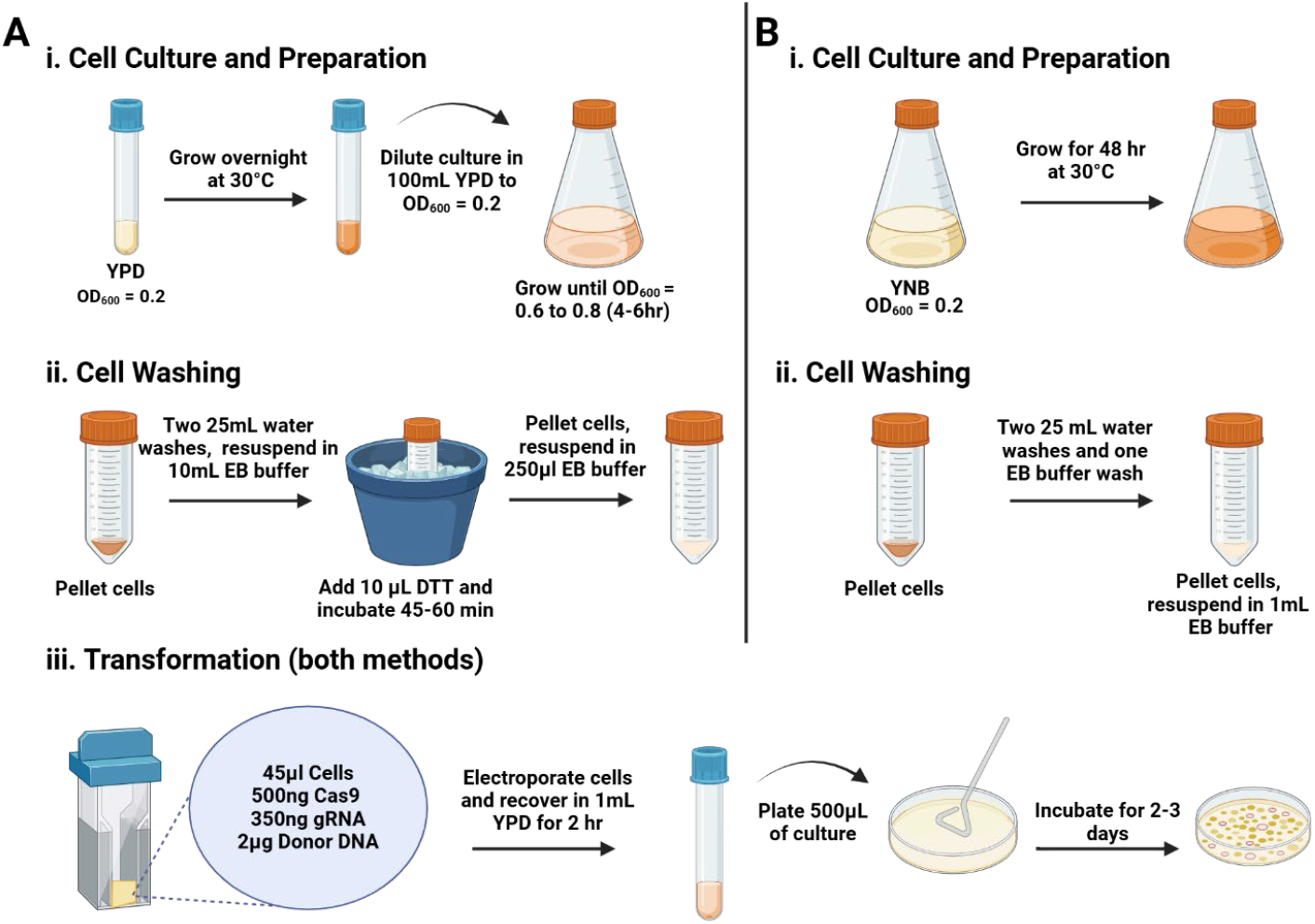
Graphical synopsis of cell culture, preparation, and transformation protocol. Brief workflow diagram depicting the timeline and key steps for preparation of (A) competent cells and (B) YNB-grown cells. Precise details may be consulted in Materials and Methods. The image was created with Biorender.

To test whether the YNB-grown cells could be used for genetic transformation, we targeted the well-characterized phosphoribosylaminoimidazole carboxylase encoding *ADE2* gene using TRACE in YNB-grown or competent cells. Deletion of *ADE2* is a common way to test gene deletion systems, as loss of this gene results in the easily observable phenotypes of red mutant colonies (5). For the transformation, 500ng of the *CnoCas9* cassette, 350ng of the sgRNA cassette, and 2mg of the gene deletion construct were mixed with YNB-grown or competent cells and electroporated. Transformation without the Cas9 cassette was also included for a CRISPR control. The transformations were plated and selected on YPD plates supplemented with nourseothricin (NAT) for transformants.

Transformants were obtained for both the YNB-grown and competent cells. The competent cells exhibited around a 65% (±4.70%) frequency of *ADE2* disruption, as indicated by the number of red colonies on the plates (Figure 1 and Table 1). This is constant with the reported *ADE2* disruption frequency by TRACE at the concentration we tested (5). Excitedly, *ADE2* disruptions were also observed with the YNB-grown cells (Figure 2). With around a 97% (±0.37%) frequency of ADE2 disruption, the transformation efficiency was significantly better than the competent cells (Figure 2 and Table 1). The YNB-grown and competent cell CRISPR control plates had transformants consisting primarily of white colonies (Figure 2). These data show that YNB-grown cells can be used directly for genetic transformation and may be more efficient than traditional competent cells.

**Table 1.** Deletion of *ADE2* with a 1-kb arm deletion construct.

| Table 1. Deletion of *ADE2* with a 1-kb arm deletion construct. |  |  |  |  |
| --- | --- | --- | --- | --- |
| Experiment | White colonies | Red colonies | Disruption frequency | Average |
| Competent | 256 | 642 | 71.49% | 64.33% |
| Competent | 434 | 541 | 55.49% |  |
| Competent | 277 | 538 | 66.01% |  |
| YNB | 83 | 1962 | 95.94% | 96.61% |
| YNB | 150 | 5238 | 97.22% |  |
| YNB | 136 | 3954 | 96.67% |  |

**Figure 2:**
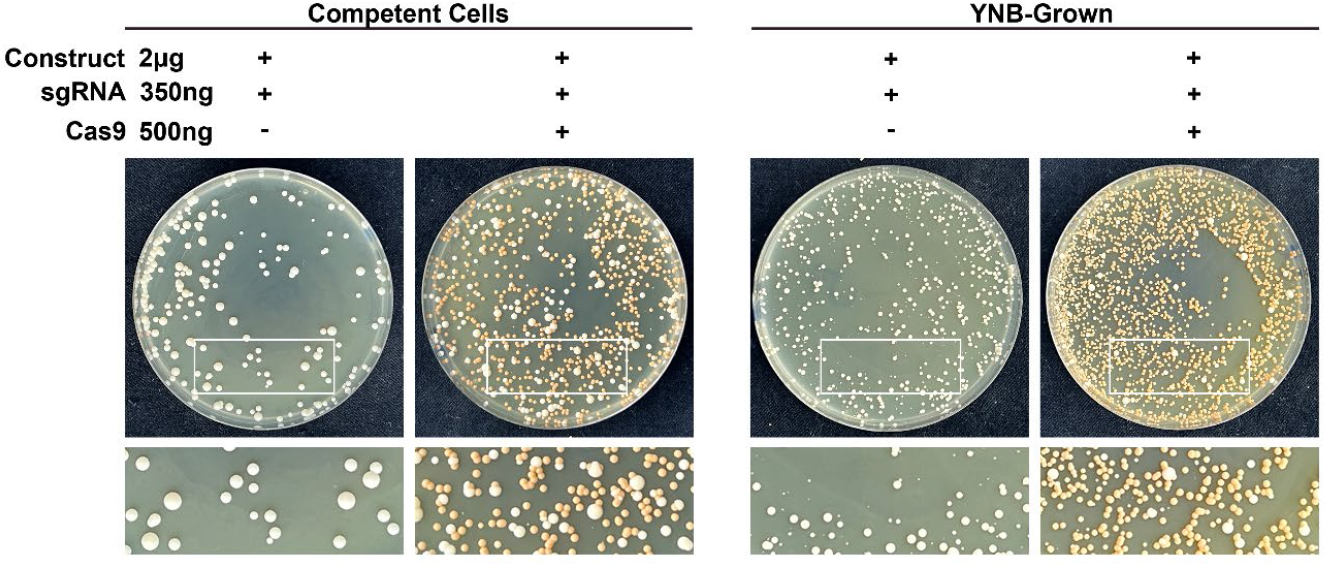
YNB-grown cells can be used directly for genetic transformation. Transformation plates for deletion of *ADE2* using competent cells or YNB-grown cells using TRACE. Transformation without the Cas9 cassette was also included for a CRISPR control. The transformations were plated and selected on YPD plates supplemented with nourseothricin (NAT).

### YNB-grown cells improve deletions using short (50 bp) homology arms

The rates of homologous replacement in *C. neoformans* are relatively low (13–15). Due to this, long homologous flanking arms (∼1 kb) are needed for targeted gene deletion in this organism. This requires multiple cloning steps or fusion PCRs to generate the deletion constructs, which adds to the technical difficulties in making gene deletion. Recently, using a *C. neoformans*-optimized Cas9, it was shown that gene deletion could be accomplished using short (50 bp) homology arms with the TRACE system (10). Since the transformation efficiency of YNB-grown cells is significantly better than the competent cells, we next sought to test if this would also be true for short homology arms.

To make the short-homology gene deletion construct, 50-bp sequences for the *ADE2* homologous arms were directly included in the primers, and the *ADE2* gene deletion construct was made by a single round of PCR from a plasmid carrying the NAT drug marker. YNB-grown or competent cells were then transformed using the TRACE system as described above. Once again, both the YNB-grown and competent cell CRISPR control plates had transformants consisting primarily of white colonies (Figure 3). The competent cells exhibited around a 44% (±5.04%) frequency of ADE2 disruption, as indicated by the number of red colonies on the plates (Figure 3 and Table 2). This was less efficient than the long homology arms seen above (Figure 2 and Table 1). Once again, YNB-grown cells had a significantly better transformation efficiency than the competent cells, with around a 93% (±2.14%) frequency of ADE2 disruption (Figure 3 and Table 2).

**Table 2.** Disruption of *ADE2* and *LAC1* with a short arm deletion construct.

| Table 2. Disruption of *ADE2* and *LAC1* with a short arm deletion construct. |  |  |  |  |
| --- | --- | --- | --- | --- |
| Experiment | White colonies | Red colonies | *ADE2* disruption frequency | Average |
| Competent | 291 | 337 | 53.66% | 44.13% |
| Competent | 414 | 238 | 36.50% |  |
| Competent | 599 | 438 | 42.24% |  |
| YNB | 213 | 2052 | 88.74% | 93% |
| YNB | 105 | 2030 | 94.83% |  |
| YNB | 95 | 2056 | 95.43% |  |
| Experiment | Black colonies | White colonies | *LAC1* disruption frequency | Average |
| Competent | 36 | 64 | 64% | 53% |
| Competent | 53 | 47 | 47% |  |
| Competent | 52 | 48 | 48% |  |
| YNB | 31 | 69 | 69% | 68.33% |
| YNB | 33 | 67 | 67% |  |
| YNB | 31 | 69 | 69% |  |

**Figure 3:**
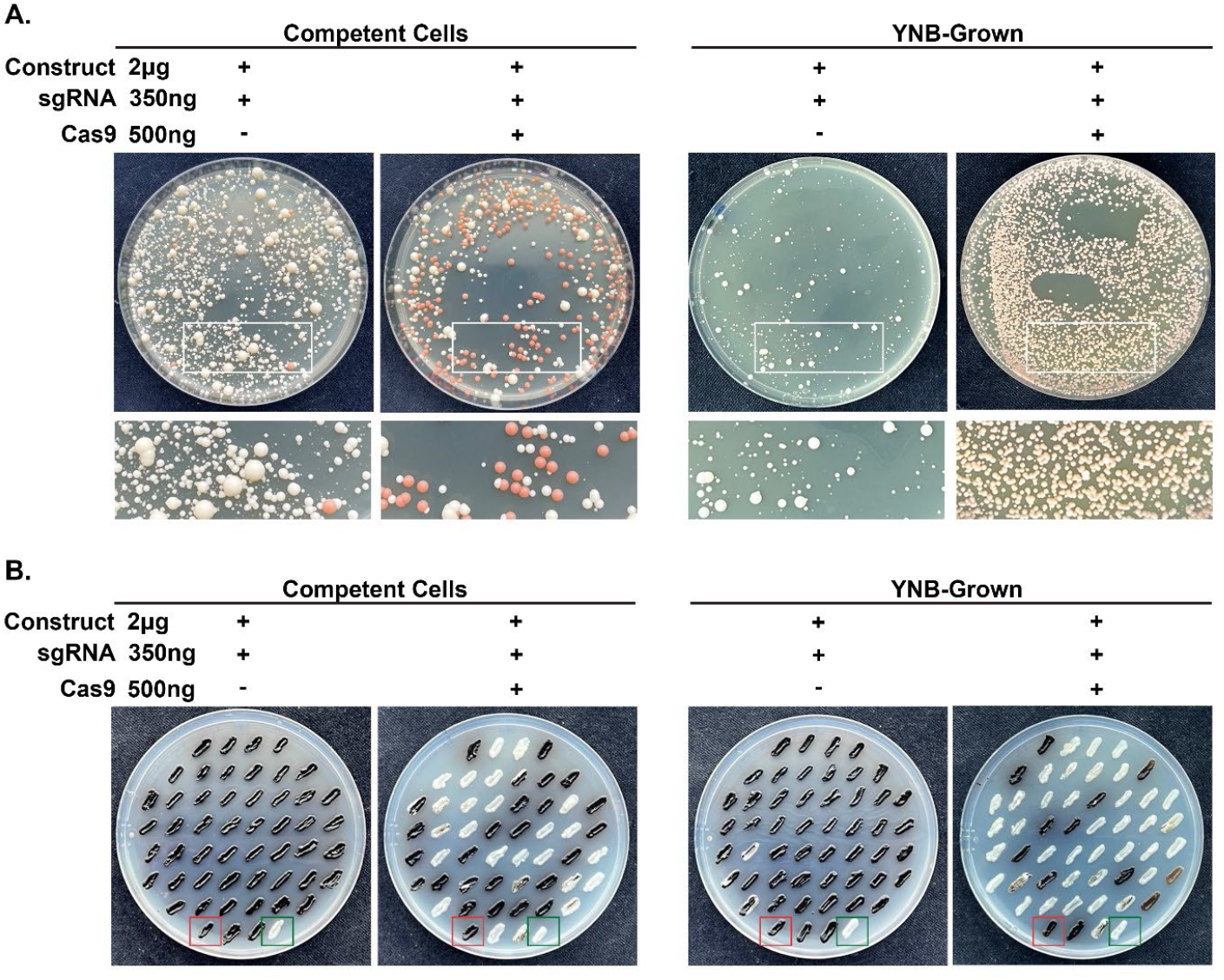
YNB-grown cells improve deletions using short (50 bp) homology arms. (A) Transformation plates for deletion of *ADE2* using competent cells or YNB-grown cells using short homology deletion cassette. Transformation without the Cas9 cassette was also included for a CRISPR control. The transformations were plated and selected on YPD plates supplemented with nourseothricin (NAT). (B) Image of a *LAC1* deletion transformation using short homology deletion cassette plate patched onto L-DOPA plates. The wild-type (WT) parental strain (red box) and a known confirmed *lac1Δ* strain (green box) were also added to each plate for controls.

To confirm our findings, we chose to delete a different gene. The laccase gene (*LAC1*) is involved in producing 3,4-dihydroxyphenylalanine (DOPA)-melanin in *C. neoformans* (16). Mutants lacking *LAC1* cannot make melanin and appear white instead of black when plated on L-DOPA media, allowing easy identification of *LAC1* disruption. To make the gene deletion construct, 50-bp sequences for the *LAC1* homologous arms were directly included in the primers, and the *LAC1* gene deletion construct was made by a single round of PCR from a plasmid carrying the NAT drug marker. YNB-grown or competent cells were then transformed using the TRACE system as described above. To screen for *LAC1* disruption, 100 colonies were randomly selected from each plate and patched onto L-DOPA plates. The wild-type (WT) parental strain and a known confirmed *lac1Δ* strain were also added to each plate for controls. The YNB-grown and competent cell CRISPR control plates had transformants consisting primarily of WT black colonies (Figure 3). While not as big a difference as seen with ADE2, YNB-grown cells also had better transformation efficiency than the competent cells, with around 68% (±0.67%) for YNB-grown and around 53% (±5.51%) for the competent cells (Figure 3 and Table 2).

### Short arm homology-directed repair is significantly improved in YNB-grown cells

YNB-grown cells had significantly better transformation efficiency than competent cells using short (50 bp) homology arms with the TRACE system. However, we do not know if the *ADE2* or *LAC1* genes were deleted or disrupted leading to the change in color. The group that developed TRACE tried using short homology arms. However, they found that this led to the insertion of the drug marker into the target locus instead of gene deletion (5). Using a transiently expressed codon-optimized CAS9, a different group was able to make gene deletions using short (50 bp) homology arms. Still, like the first paper, they found it inefficient, leading to more insertions and gene disruption than gene deletions (10). To test the deletion efficiency YNB-grown cells, we devised a PCR screen. To do this, 10 red colonies from the *ADE2* transformation plates or 10 white colonies from the L-DOPA plates were randomly selected from the YNB-grown or competent cell groups, and genomic DNA was isolated from each mutant. Homology-directed repair (HDR) at the 5′ or 3′ end of the genes was assessed using a three-primer strategy. Three primers were used to ensure that there would be a product corresponding to the WT or mutant band. Strains with mutant bands at both the gene’s 5′ and 3′ ends were considered to have the gene deleted. Mutants of *ADE2* or *LAC1* from the competent cell were mainly gene disruptions, with only 6.7% of the tested mutants with HDR at both ends (Figure 4). This is constant with the reported use of short homology arms (10). Excitedly, YNB-grown cells had a significantly higher number of gene deletions for both *ADE2* (63.3%) and *LAC1* (56.7%) than the competent cells (Figure 4). These data show that short arm HDR is significantly improved in YNB-grown cells.

**Figure 4:**
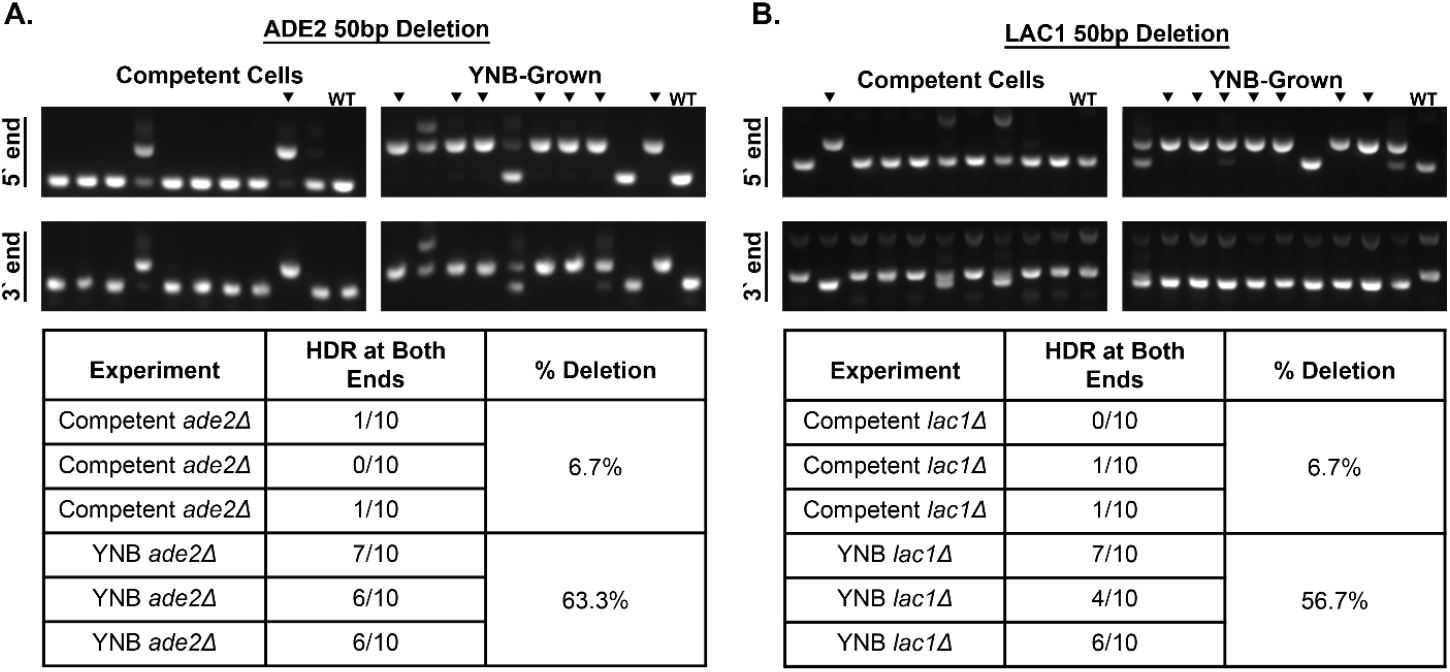
Short arm homology-directed repair is significantly improved in YNB-grown cells. Genotyping of (A) *ADE2* and (B) *LAC1* transformants. Representative gels from PCR genotyping of transformants. Ten red colonies from the *ADE2* plates or ten white colonies from the *LAC1* plates were randomly selected from each condition and screened. Black arrowheads indicate transformants with HDR at both ends. Data was collected from 3 independent experiments.

## Discussion

The fungal cell wall is a significant barrier to successful fungal transformations, as exogenous DNA must first pass through it. Most transformation protocols have steps that remove or damage the cell wall. Standard transformation protocols for most filamentous fungi and some yeast involve the formation of protoplasts by treatment with cell wall-degrading enzymes to remove the cell wall. Treatment with dithiothreitol (DTT) can improve the competency of the fungal cells by the reduction of cell wall proteins and a general increase in cell wall porosity by reducing disulfide bridges (6, 17). Transformation efficiency can also be improved by changing the concentrations of DTT or by adding additional chemicals like lithium acetate (18). Additionally, fungal mutants that have altered cell wall structure have increased transformation efficiency. During a mutant screen in *Saccharomyces cerevisiae, pde2Δ, pmr1Δ*, and *spf1Δ* mutants were identified as highly transformable (19). These three mutants all have changes in cell wall structures. Further studies with the *spf1Δ* mutant using a cell-impermeable fluorescent DNA probe showed that the increase in transformability was most likely due to more exogenous DNA passing through the cell wall and entering the cell during transformation (19). This is likely the reason why the YNB-grown cells have increased transformation efficiency, as portions of the capsule and outer wall are stripped away from the cells, leading to a significantly altered cell wall structure.

In addition to improving transformation efficiency with constructs with long homology arms, we also found that YNB-grown cells improve gene deletions with constructs with short homology arms (Figure 4). The use of short homology arms for genetic manipulations in *C. neoformans* leads primarily to gene disruptions over gene deletions. Recently, it was reported that using a cryptococcal optimized Cas9 improved the efficiency of short (50 bp) homology arm constructs with the TRACE system (10). Efficiency was further improved when the optimized Cas9 was stably integrated and constitutively expressed (10). In this system, the optimized Cas9 was integrated into the Safe Haven 2 site (10). The authors found that they could significantly increase the number of *URA5* deletion mutants over insertion mutants in the Cas9 integrated strain compared to when Cas9 was transiently expressed (10). While this has significantly improved the ability to use constructs with short homology arms, a disadvantage of the system is that you need to insert Cas9 into the background strain you are using. In our system using simple growth conditions, we are also able to dramatically improve the number of deletion mutants over insertion mutants (Figure 4) using the optimized Cas9. An advantage of our system is that we can use unmodified strains to make deletions with improved efficiency.

This study found that growing cryptococcal cells in unbuffered YNB media for 48 hours can be used directly for genetic transformation. YNB-grown cells are more efficient than traditional competent cells with both long and short-homology deletion constructs. When using short-homology deletion constructs, YNB-grown cells have a greater rate of deletion mutants over insertion mutants. Using simple growth conditions, we have greatly improved the speed and efficiency of cryptococcal genetic transformations.

## Materials and Methods

### Strains and media

*C. neoformans* strains KN99-α and *lac1Δ* were a gift from Jennifer Lodge. Strains were stored at −80°C as glycerol stocks. Strains were revived and maintained on YPD (1% yeast extract, 2% Bacto peptone, and 2% dextrose) solid medium containing 2% Bacto agar. Cryptococcal transformants were selected on YPD medium supplemented with 100 μg/ml nourseothricin (NAT) (Werner BioAgents).

### Generation of the Cas9, sgRNA, and the gene deletion constructs

The Cas9 (pBHM2403) and sgRNA (pBHM2329) plasmids were a gift from Hiten Madhani (Addgene plasmids # 171687 and # 171686) and the NAT (pGMC200) plasmid was a gift from Jennifer Lodge. All primers were ordered from IDT (see Table S1 for primer list)

The CAS9 cassette was amplified from pBHM2403 using Phusion Flash High-Fidelity PCR Master Mix (ThermoFisher, #F548L) per the manufacturer’s instructions and primers Cas9_P1 and Cas9_P2.

The *ADE2* (CNAG_02294) and *LAC1* (CNAG_03465) sgRNA sequences were designed using EuPaGDT (http://grna.ctegd.uga.edu/). To make the sgRNA construct, the U6 promoter was amplified with the 20 bp target sequence from pBHM2329 using gRNA_P1 and gRNA_P2_ADE2 or gRNA_P2_LAC1 primers. The 20 bp target sequence with the sgRNA scaffold and 6T was then amplified from pBHM2329 using gRNA_P3_ADE2 or gRNA_P3_LAC1 and gRNA_P4. The two products were then joined by fusion PCR using primers gRNA_P5 and gRNA_P6.

The long arm gene deletion construct for *ADE2* was generated using overlap PCR. Briefly, the 5′arm upstream of the start site and the 3′ arm downstream of the stop site were amplified from KN99 genomic DNA using primers ADE2_P1 and ADE2_P4 or ADE2_P2 and ADE2_P5. The NAT resistance cassette was amplified from pGMC200 using primers ADE2_P3 and ADE2_P6. The three products were then joined by fusion PCR using primers ADE2_P7 and ADE2_P8. The short homology arm deletion constructs for *ADE2* and *LAC1* were amplified from pGMC200 using primers ADE2_F_50 and ADE2_R_50 or LAC1_F_50 and LAC1_R_50. The sizes of PCR products were verified by gel electrophoresis and all PCR products were cleaned and concentrated using the PureLink PCR purification kit (Invitrogen, #K310001) and eluted in sterile ddH2O.

### Preparation of electrocompetent cells

Electrocompetent cells were prepared as described by Lin and colleagues (20). Briefly, overnight KN99 cultures were back diluted to an OD600 of 0.2 into 100ml of YPD and grown until the culture reached the mid-log phase (OD600 of 0.6-0.8). The cells were then washed with ice cold sterile ddH2O twice and then resuspended in 10 ml of electroporation buffer (10 mM Tris-HCl pH 7.5, 1 mM MgCl2, 270 mM sucrose). Next, 10µl of 1M dithiothreitol (DTT) was added to the cells and then incubated on ice for one hour. The cells were washed with electroporation buffer to remove the DTT, resuspended in 250µl of electroporation buffer, and kept on ice.

For YNB-grown cell, a 250ml flask containing 50ml of unbuffered Yeast Nitrogen Base (YNB) media (0.67% yeast nitrogen base without amino acids [Difco #291940] containing 2% dextrose) was inoculated with KN99 at an OD600 of 0.2 and was grown at 30°C in a shaking incubator at 300 rpm for 48 h. After 48h, the cells were washed with ice-cold sterile ddH2O twice and then once in electroporation buffer, resuspended in 1ml of electroporation buffer, and kept on ice.

### Electroporation

The cells were transformed by electroporation. Briefly, 45ul of either the competent or YNB-grown cells were mixed with 5 µl of DNA mix containing 2 µg deletion construct, 350 ng sgRNA, and 500 ng CAS9, or 2 µg deletion construct and 350 ng sgRNA with no CAS9 as a control. The cell mix was then transferred to a precooled 2 mm gap electroporation cuvette (Bio-Rad Laboratories). A Gene Pulser Xcell Total System (Bio-Rad Laboratories) was used for electroporation with the following settings: 450 V, 400 Ω, 250 µF. Cells were resuspended following electroporation in 1 ml YPD and incubated at 30°C for 2 h before plating on selective media.

### Phenotyping

To test for *ADE2* disruption, the recovered cells were plated onto selective media and incubated for three days at 30°C. After three days, the plates were moved to 4°C for one week to allow stronger red pigmentation to develop and then imaged. To test for *LAC1* disruption, the recovered cells were plated onto selective media and incubated for three days at 30°C. After three days, 100 colonies were randomly selected and patched onto YPD and L-DOPA plates (13 mM glycine, 15 mM glucose, 29.4 mM KH2PO4, 10 mM MgSO4 ⋅ 7H2O, 3 µM thiamine, 5 mM D-biotin, 1 mM L-3,4-dihydroxyphenylalanine [L-DOPA], and 2% agar) for three days at 30°C in the dark. The wild-type (WT) parental strain and a known confirmed *lac1Δ* strain were also added to each plate for controls. After three days, the plates were imaged.

### PCR genotyping

To test if the *ADE2* or *LAC1* genes were deleted or disrupted, the mutants were screened by PCR. To do this, 10 red colonies from the *ADE2* plates or 10 white colonies from the *LAC1* plates were randomly selected, and genomic DNA was isolated from each mutant using the Monarch Genomic DNA Purification Kit (New England Biolabs). The three primers ADE2_check1, ADE2_check1, and NAT_check_5′ were used to check the 5 prime *ADE2* locus, and the three primers ADE2_check3, ADE2_check4 and NAT_check_3′ were used to check the 3 prime *ADE2* locus. The three primers LAC1_check1, LAC1_check1, and NAT_check_5′ were used to check the 5 prime *LAC1* locus and the three primers LAC1_check3, LAC1_check4, and NAT_check_3′ were used to check the 3 prime *LAC1* locus. Three primers were used to ensure that there would be a product corresponding to the WT or mutant band.

## Acknowledgments

This work was supported by a National Institutes of Health grant K22 AI148724, the Center of Pediatric Experimental Therapeutics, and start-up funds from the Department of Clinical Pharmacy and Translational Science. The funders had no role in study design, data collection, and interpretation or the decision to submit the work for publication.

---

**Supplemental Table 1.**
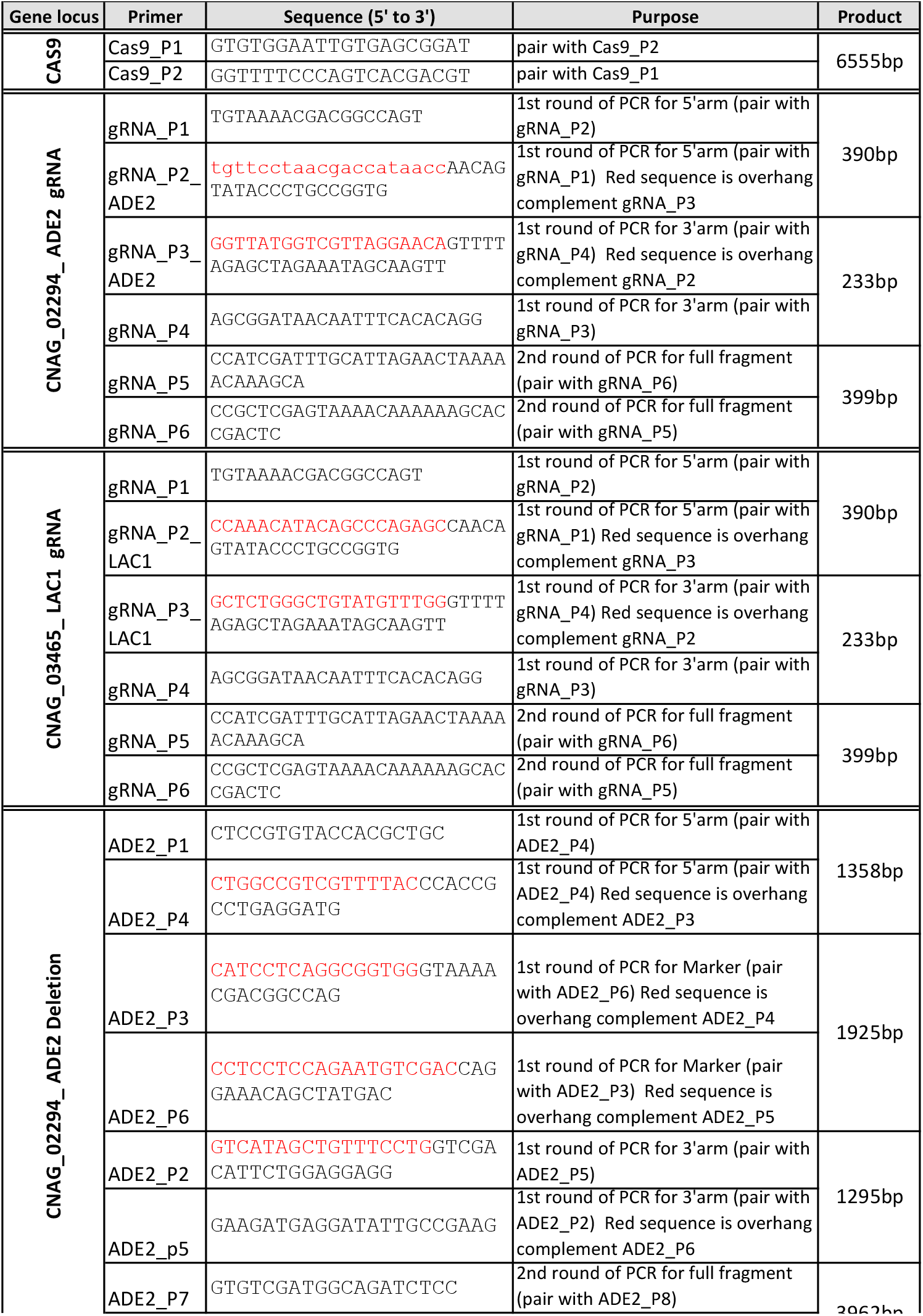
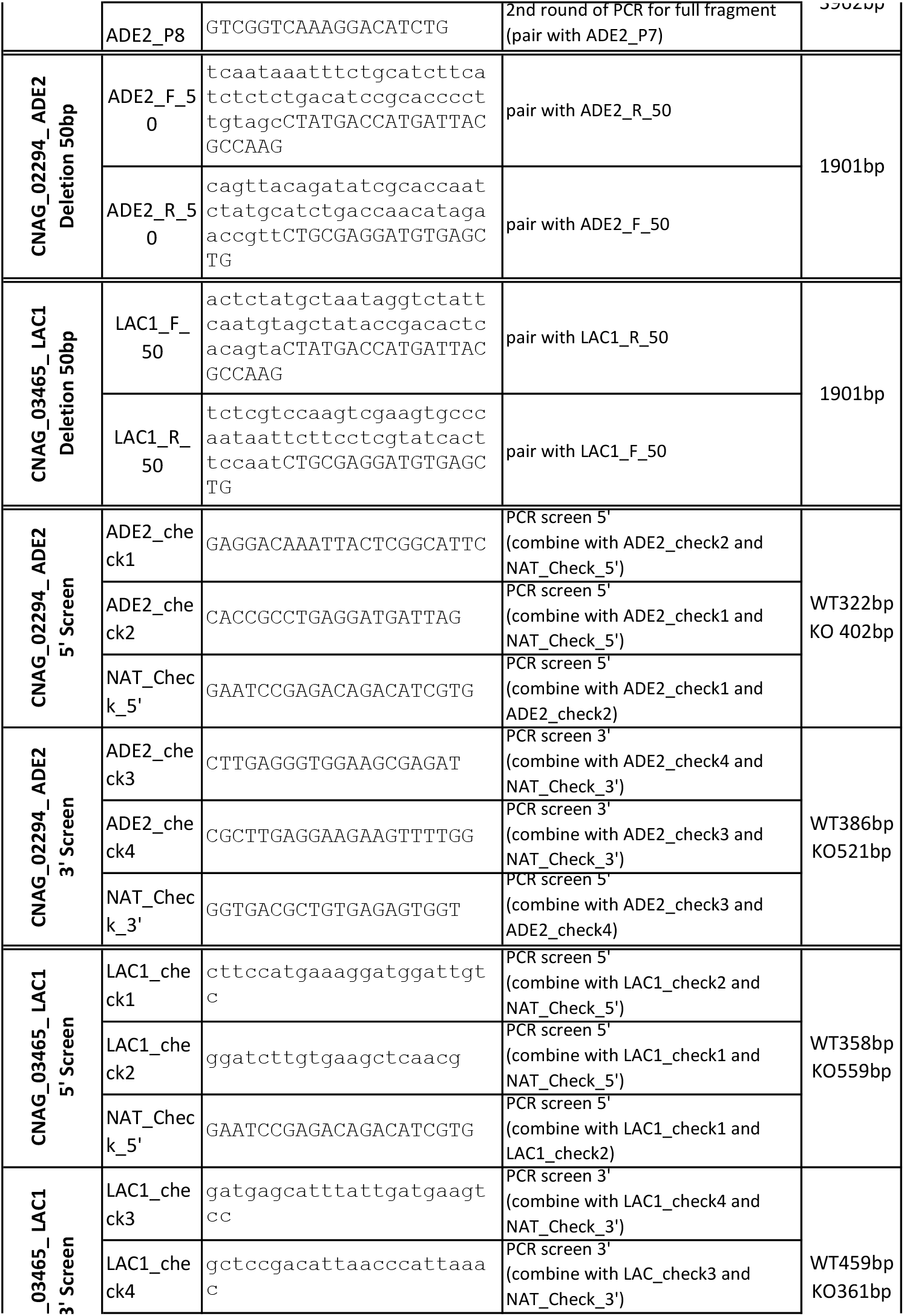
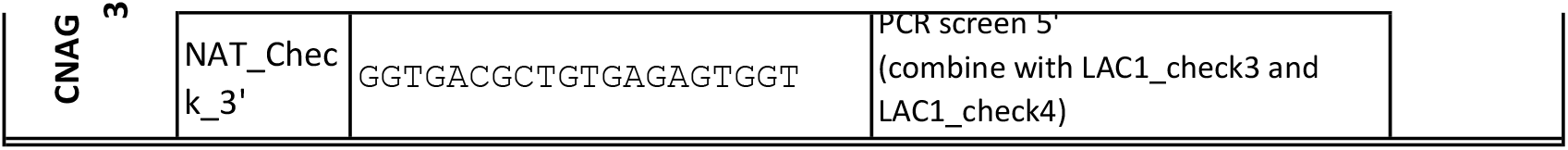
Primer Used In Study.

